# Integrative Taxonomy: From FASTA Pain to Species Gain with the interactive “IntegraTax”

**DOI:** 10.64898/2025.12.22.695429

**Authors:** Amrita Srivathsan, Leshon Lee, Vivian Feng, Rudolf Meier

## Abstract

Here we present IntegraTax, a tool for analysing and managing taxonomic projects that combine DNA data with other evidence such as morphology to arrive at integrative species boundaries. IntegraTax visualizes genetic clusters through single-linkage clustering (“Objective Clustering”) and provides an interactive browser interface that allows users to record taxonomic decisions regarding species limits. Projects can be saved at any stage, thus allowing continuous tracking of annotations and taxonomic decisions across many sessions. A typical IntegraTax session starts with a set of sequences that are visualized as a cluster fusion diagram revealing the genetic distances between the sequences and clusters. Users can define an “instability zone” to distinguish clusters that are clearly distinct, likely representing separate species, from those whose status is uncertain based on genetic data alone. Based on the instability zone setting, IntegraTax then suggests which and how many specimens should be studied with a second source of data to validate species hypotheses. This process is facilitated by an interactive html environment that enables detailed specimen-level annotations. For example, a taxonomist can label which specimens have been studied, which clusters have been validated as species, and which species can be identified. By combining clustering, intuitive visualization, and easy annotation in one interactive framework, IntegraTax treats species hypotheses as annotated objects that can be inspected, revised, and exported at any stage, with documentation of examined specimens. It simultaneously allows researchers to manage taxonomy projects with tens of thousands of specimens and hundreds of species. This will become increasingly important as taxonomists start resolving the species boundaries of the millions of undescribed species particularly within hyperdiverse dark taxa.

## Introduction

Taxonomy faces many long-standing problems. Two interconnected challenges are that most species remain undescribed, a pattern consistently demonstrated for hyperdiverse taxa where undescribed diversity regularly exceeds named species by factors of ten or more (“dark taxa”) (Hartop et al., 2022). The second problem is that approximately five million names exist for only <2.5 million valid species (Bánki et al., 2025, GBIF, 2025). This redundancy is caused by inadequate species descriptions, inaccessible literature, hidden/lost types, and unstable species boundaries. This is all bad, but there is also good news. Perhaps only 20% of all species have been described so far (Mora et al., 2011). This means that we can do better for the remaining 80%. One way to do better is to embrace integrative taxonomy (Dayrat, 2005, Padial et al., 2010). This is because species boundaries are likely more stable when they are determined based on multiple data sources (Dayrat, 2005, Padial et al., 2010). Furthermore, identification of specimens to species will also be less likely to be incorrect when different kinds of data can be used. Overall, integrative taxonomy should thus lower the risk of producing synonyms while also reducing how often species have to be split or merged. Most taxonomists thus agree that integrative taxonomy is the way forward (Dayrat, 2005).

However, collecting and analysing more than one type of data for all specimens also reduces the number of species that can be described within a given amount of time. Every additional data source may improve the stability of species boundaries and the ability to identify specimens, but it also slows the overall discovery process. To mitigate these downsides, large-scale integrative studies have to be efficient. One way to achieve efficiency is through a two-step process in which one type of data that is easily generated is used to generate preliminary species hypotheses, while a second and more inaccessible source of evidence is only acquired for selected few specimens to test these hypotheses (Hartop et al., 2022). Historically, this workflow began with morphospecies sorting (often by parataxonomists) followed by barcoding of a few specimens for each putative species. However, this workflow required so much manpower for large samples that many dark taxa were never sorted (Meier et al., 2016, Wang et al., 2018). With the increasing cost-effectiveness of DNA sequencing, the two-step process has reversed (Hebert et al., 2018, Meier et al., 2016, Srivathsan et al., 2021). DNA barcodes are now used to generate initial species hypotheses which are then tested with morphology (Wang et al., 2018). This “reverse workflow” is now central to many large-scale integrative and dark-taxon studies (Caruso et al., 2024, Hébert and Favret, 2025, Meier et al., 2025b, Hartop et al., 2022, Wang et al., 2018, Srivathsan et al., 2019) because it reduces the number of specimens that require detailed morphological examination. Furthermore, several empirical studies have shown that picking specimens for morphological investigation based on haplotype information is very effective at finding and resolving conflict between DNA barcode clusters and morphospecies (Amorim et al., 2025, Caruso et al., 2024, Gustafsson et al., 2025, Hartop et al., 2022, Hébert and Favret, 2025, Lee et al., 2025, Meier et al., 2025b, Riccardi and Hartop, 2024).

However, even this streamlined two-step approach still requires dedicated tools for managing the process, because it is not trivial to generate species hypotheses, track which specimens have been examined, evaluate supporting evidence, and record taxonomic decisions that may change over time. Currently, such tracking is still handled with spreadsheets or even handwritten notes, but neither scale, and both are likely to cause mistakes when thousands of specimens belonging to hundreds of species are studied. This is one major motivation for developing IntegraTax. It takes a set of sequences, generates hierarchical barcode clusters and then allows the user to determine which clusters correspond to morphospecies through the study of a subset of specimens with morphology. The results of the examination are saved in an interactive environment. To facilitate the process, IntegraTax even suggests which specimens should be studied with morphology by implementing the rules developed in an empirical study on a dark taxon (Hartop et al., 2022). Taxonomic decisions can be made across many sessions, and one can create a version history by regularly saving working files under different names.

### The two modules of IntegraTax

IntegraTax contains two modules and is packaged as a combination of the two (https://github.com/asrivathsan/IntegraTax/releases). Module two can be separately downloaded.

#### Module 1: Objective Clustering

To start an integrative taxonomy project, the “clustering” module of IntegraTax is used to create a hierarchical cluster diagram based on barcode data. The software uses single-linkage clustering (“Objective Clustering”: Meier et al., 2006) because it allows for showing the fusion points as pairwise-distances (p-distances) on the nodes of the diagram. Cluster statistics reveal how many clusters have been found at all fusion points (e.g., at 1%, 2%, 3% p-distance). Below, we motivate the methodological choices which we believe serve taxonomists well because they avoid unnecessary assumptions and transformations.

IntegraTax implements clustering with p-distances instead of Kimura Two-Parameter (K2P) distances because model testing always confirms that K2P is not an appropriate model for *Cytochrome Oxidase I* (*COI*) data (Collins et al., 2012, Meier et al., 2006, Srivathsan and Meier, 2012). This is not surprising because closely related sequences are unlikely to be affected by hidden substitutions through multiple hits, making model corrections unnecessary. Furthermore, correcting for transition-transversion biases with K2P is not likely to yield satisfying estimates, given that K2P models the data under the assumption of equal base frequencies, uniform substitution processes, independence across sites and lineages based on very short barcode fragments that exhibit compositional biases and often code for proteins (Pentinsaari et al., 2016). An additional reason to prefer p-distances over K2P-distances is that the latter widens barcode gaps when interspecific distances are larger than intraspecific distances. This carries the risk of creating systematic biases by producing larger numbers and more sharply separated clusters than implied by the data.

IntegraTax uses single-linkage clustering for visualizing barcode data instead of neighbour joining trees, because the latter require the conversion of a pairwise distance matrix into bifurcating trees by estimating branch lengths that minimize the total distortion between observed and the implied distances on the tree. When many sequences are identical or near identical, a large number of trees have equal fit (“tie trees”) which is then resolved by arbitrarily picking one of the bifurcating trees (Takezaki, 1998). The effect of this arbitrariness is subsequently reduced by producing a consensus tree before or after running a bootstrap analysis. It is not clear how this procedure could help with representing the barcode data in an integrative taxonomy study and its use has been repeatedly criticized in literature (Collins and Cruickshank, 2013, Will and Rubinoff, 2004).

Single-linkage clustering yields a unique solution for a given set of barcodes and directly indicates the pairwise distances at which barcode clusters merge. Its use in DNA barcoding has a long history. Early implementations, such as that described by Blaxter et al. (2005), relied on order-dependent procedures, whereas “objective clustering” in SpeciesIdentifier presented a reproducible formulation based on distance connectivity (Meier et al., 2006). Not surprisingly, single-linkage clustering thus also underpins many species delimitation algorithms. For example, Automatic Barcode Gap Discovery (ABGD; Puillandre et al., 2012) and Assemble Species by Automatic Partitioning (ASAP; Puillandre et al., 2021) apply distance-connectivity rules equivalent to single-linkage clustering while additionally identifying barcode gaps or scoring schemes to propose delimitation hypotheses. The unpublished RESL algorithm that yields Barcode Index Numbers (BINs; Ratnasingham & Hebert, 2013) likewise appears to start from single-linkage clustering partitions. The use is motivated because numerous empirical studies revealed that threshold-based clusters have good congruence with morphology (Ahrens et al., 2016, Hartop et al., 2022, Hawlitschek et al., 2013, Meier et al., 2025b, Slater-Baker et al., 2025, Srivathsan et al., 2019, Wang et al., 2018, Vogel et al., 2024). This is particularly relevant for integrative taxonomy, because the number of specimens requiring morphological examination increases with every molecular cluster that is incongruent with morphological evidence (Hartop et al., 2022).

Single-linkage clustering, however, has one undesirable property. The distances within a cluster can exceed the fusion distance as discussed in Meier et al. (2006). Therefore, IntegraTax allows for displaying maximum intracluster distances in addition to fusion distances. It also records the specimens representing maximum distances. This information is then used by the Large Scale Integrative Taxonomy (“LIT”) workflow described later.

There are numerous other ways to generate species-level hypotheses based on barcode data (Ramirez et al., 2023). Besides ABGD, ASAP, and BINs, there are tree based approaches such as Poisson Tree Processes (PTP: Zhang et al.(2013)), Generalized Mixed Yule Coalescent (GMYC: Pons et al. (2006), Fujisawa & Barraclough (2013)). Recent efforts to standardize species delimitation outputs in the form of SPART format (Miralles et al., 2022), have enabled an easy integration of different approaches for visualization in IntegraTax. SPART outputs from methods such as ASAP/ABGD are already easily produced using tools such as SPARTexplorer (https://spartexplorer.mnhn.fr/) and the format allows alternative species hypotheses to be documented. The delimitations from alternative partitioning schemes are summarized by IntegraTax to visualize areas where the schemes are in disagreement and the extent of congruence across schemes. The only complication is that the single-linkage clustering used for visualization in IntegraTax may not fully reproduce the ordering or hierarchical structure implied by alternative partitioning methods, since these tend to rely on distance transformations, model-based trees, or stochastic processes: in such cases the discontinuous sections are visually highlighted for the user. However, our experience is that rearrangements at the species level are usually minor as the underlying barcode sequences are the same across approaches. BINs or other species delimitations can likewise be displayed on cluster fusion diagrams through FASTA header annotation or by creating a SPART-equivalent file, but we caution against the use of BINs because BIN memberships routinely change when new sequences are added to BOLD (Meier et al., 2022, Page, 2025). This complicates the use of BIN identifiers since the boundaries may have shifted between the start and end of a project. Furthermore, the use of BINs in scientific studies is problematic until the underlying clustering algorithm is published, an implementation that can be run independently is made available, and the algorithm in BOLD excludes private data for BIN calculation.

Most taxonomists carrying out an integrative taxonomy study will likely combine new data with a set of identified reference sequences and thus IntegraTax has options to allow clustering of external sequences with project sequences (Figure 1). This can, however, create complications because the data from public databases are often replete with unidentified barcodes. This slows clustering and renders the output unwieldy for taxonomic work. Therefore, users have the option to restrict the import of external data to only sequences with binomial names and/or barcodes that fall within a specified similarity window of the project barcodes (default: 5%). In both cases, joint clustering of project and external sequences starts after a two-file upload in a GUI followed by user-specified filtering of the external data if desired. The inclusion of external sequences is useful because it helps with cluster identification when barcode clusters are composed of project sequences and identified external sequences. It also helps with determining species boundaries, for example, when external sequences show that an apparent barcode gap in the project is an artifact of sampling. Once such reference-enriched datasets have been assembled and clustered, summary statistics are available to calculate match ratios, specimen congruence and related diagnostics at different thresholds (Ahrens et al., 2016, Meier et al., 2025a, Yeo et al., 2020).

**Figure 1.**
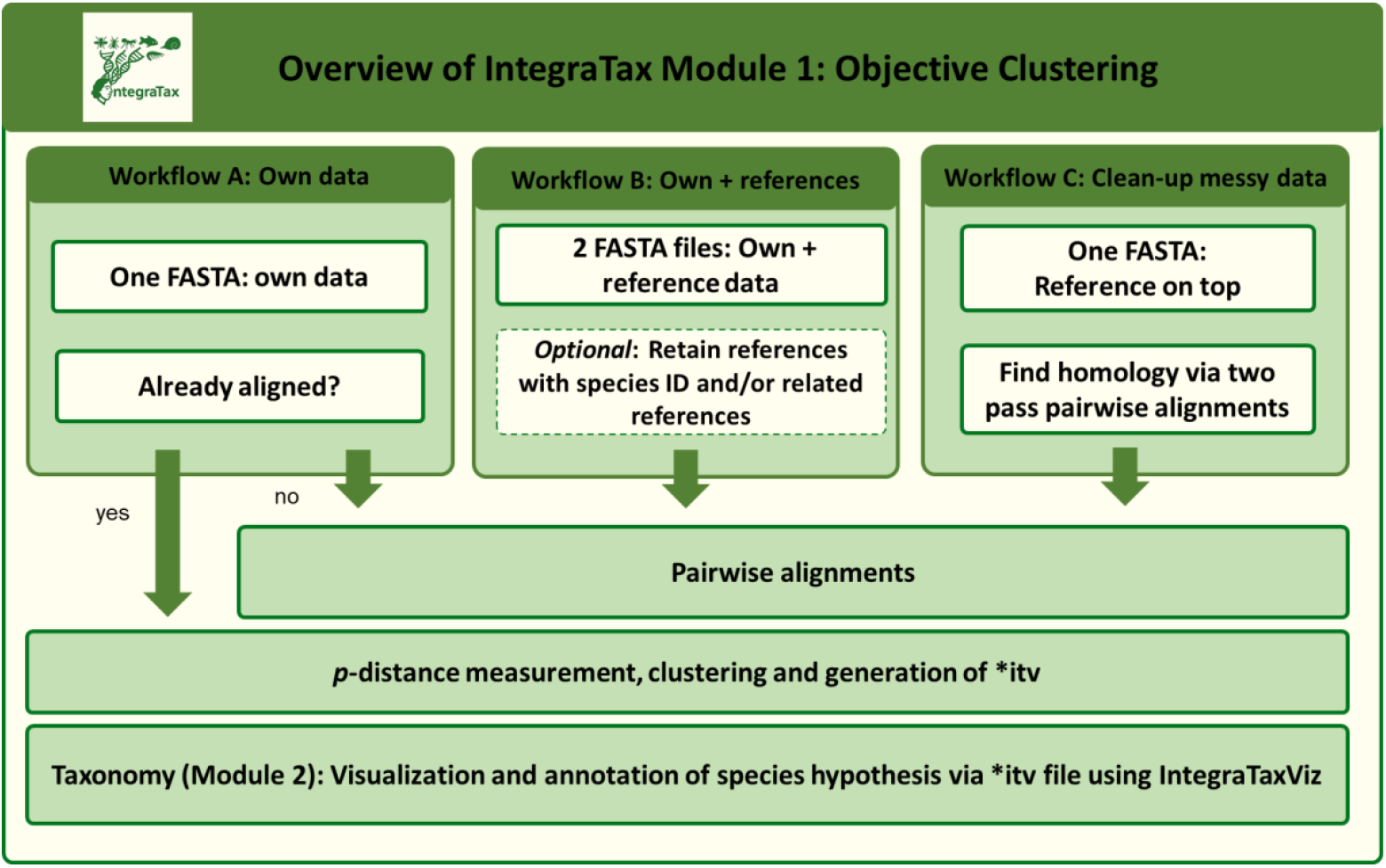
An overview of the IntegraTax clustering module showing three alternative data entry modes corresponding to common taxonomic use cases: clustering of own data only, own data with external reference data, and clean-up of heterogeneous or non-overlapping sequence datasets. Pre-aligned input is accepted only for user-provided datasets, while reference-based and heterogeneous workflows perform internal pairwise alignment. Resulting sequence clusters are inferred using p-distance and exported as an .itv file for visualization and annotation.

#### Implementation

Aligned datasets are processed through the calculation of uncorrected pairwise genetic distances. For unaligned datasets, pairwise alignments are computed using the *parasail* library (Daily, 2016), which applies Smith-Waterman alignment with affine gap penalties. Clustering in IntegraTax is implemented such that key features relevant to large-scale integrative taxonomy are simultaneously recorded. The details of implementation are as follows: A set of non-redundant sequences is first built to minimize calculations. Pairwise distance computation is fully parallelized and is conducted in batches. Distances are quantized into bins at 0.01% resolution and written as streamed batches, avoiding creation of the full pairwise distance matrix and allowing distance pairs to be processed incrementally. Distance bins are merged across the batches producing unified distance bins containing all unique sequence-pairs that fall within the distance bin.

These distance bins are then traversed in increasing order. For each bin, sequence pairs are read sequentially and used to perform single-linkage clustering: whenever a pair connects two previously distinct clusters, those clusters are merged at the current distance threshold. If a pairwise distance is encountered between sequences that already belong to the same cluster, the maximum intracluster distance for that cluster is updated accordingly. As the traversal is from smallest to largest distances, it ensures that smaller distances are never encountered later. The pair of specimens responsible for this update is recorded. This single-pass traversal over sorted distance bins ensures that clustering follows the exact distance order while remaining memory-efficient and scalable to large datasets.

This module is implemented as a software package for Windows, Mac, and Linux (Ubuntu). The largest dataset tested so far involves >50K aligned sequences in a standard MacBook Pro and clustering along with specimen assignments to clusters could be done in under 2 hours with 4 cores, while routine datasets of 10K sequences are clustered in minutes (aligned, <1 min in MacBook, 4 cores). The main limitation in terms of dataset size for IntegraTax is the visualization component (module 2) which depends on browser capacity. This has been tested for dataset sizes of up to 30K sequences. We note that visualizations are useful for integrative taxonomy when a taxonomist is unlikely to be looking at specimen sets beyond this scale simultaneously. The clustering module independently provides a text output for clustering results at every threshold where specimen grouping is changing. For threshold-based studies of large datasets, this “clusterlist” output can be directly used (Santos et al., 2025, Srivathsan et al., 2023).

Critically, this implementation enables tracking the maximum pairwise distances among a set of specimens when they fuse by the criterion of minimum distance, as well as identifying the specific specimen pairs that define the maximum distances. Thus, every cluster at every threshold is associated with three properties (minimum distance, maximum distance, and pair of specimens (haplotype representatives) that define the maximum distance) which allows users to explore data dynamically downstream. The results of the single-linkage clustering are translated into a dendrogram that forms part of the *IntegraTaxViz* input. In addition, the software allows the option to detect binomial names if they are present in the dataset or alternatively allows users to supply them as an external tab-delimited file. These names are then available within the visualization interface and can be modified for taxonomic work. The main output of the clustering module is an *IntegraTaxViz* (*.itv) file that can be opened in browsers and connects raw sequence data with the visualization of genetic distances.

Accommodating publicly available barcode data requires additional processing because external data sets tend to be large and contain sequences of different lengths. This is why IntegraTax allows the user to subset sequences based on binomial names or similarity to other barcodes. The latter is implemented by using BLAST to retrieve only related sequences from the reference FASTA file with the *blastn* default set to a 95% distance cut-off and retaining the top 100 hits per reference sequence (parameters can be modified by user). BLAST ensures both homology between sequences and reduction in dataset size. However, in the event that the user wants unrelated sequences to be retained, it is essential to protect against sets of non-overlapping sequences in the external data. IntegraTax therefore also has an exhaustive homology search option that uses the first sequence in a file as the reference region and then implements two rounds of distance calculations. The first round aligns the reference sequence to all other sequences and calculates distances for each pairwise comparison. This is implemented using the *edlib* library where edit distances are returned and the library was chosen over *parasail* due to a more comprehensive traceback option (Sosic and Sikic, 2017). The second round begins with N sequences (default = 10) sampled across the distance profile obtained from the first round and uses sequences that cover the entire reference region as seeds for a second set of N-versus-all alignments. Distance profiles composed of a mixed set of markers can usually be identified based on a breakpoint in the distance distribution.

IntegraTax uses a kernel density estimation with the widely used Silverman’s rule to detect this breakpoint. Experimental work suggests that this is effective against identifying spurious dips caused by taxonomic variation because in tests with *COI* data from Sepsidae (Insecta: Diptera) and Mycetophilidae (Insecta: Diptera), the module was able to recover 99% of the expected sequences when compared with manual curation. If automated detection is not desired, users may specify fixed distance thresholds if the breakpoints are not clear and/or if they prefer to control by specific thresholds. The output excludes non-target sequences and trims retained sequences to the expected region defined by the reference, while discarded sequences and distance profiles are retained for inspection should the user wish to adjust parameters. The excluded sequences are provided in a file, and this file should be reviewed by the user before finalizing their dataset. This is particularly important when this option is used to clean-up a project file.

#### Module 2: Taxonomy

The second module of IntegraTax is the taxonomy module. It consists of an interactive tool for annotating clusters, tracking taxonomic decisions, and identifying which specimens should be examined to confirm or revise preliminary hypotheses. Manual tracking of these decisions is difficult in large datasets, when thousands of specimens must be evaluated and manual record-keeping thus becomes unreliable. One goal of IntegraTax is to make the process explicit, repeatable, and well documented. To support this goal, the taxonomy module allows users to export curated results at any stage of the workflow, ensuring that annotations, decisions, and species hypotheses can be transferred to downstream analyses and external tools (SVG, FASTA, Excel LIT, SPART). Exports are also designed to facilitate integration with external workflows and biodiversity data portals.

Large-scale integrative taxonomy requires a structured procedure for deciding which specimens should be examined and how conflicts between molecular clusters and morphology should be resolved. As a default, IntegraTax implements the “Large-Scale Integrative Taxonomy” (LIT) criteria for specimen selection developed in an empirical study by (Hartop et al., 2022). Note that these criteria can be adapted or disregarded depending on project-specific needs. For example, LIT relies heavily on haplotype information for subsampling, but if the sharing of identical haplotypes by cryptic species is suspected to be widespread, subsampling should be extended across the geographic range of clusters. Users can furthermore change the default settings for the instability zone and intracluster variability. This will reduce or increase the number of specimens that are sampled for each cluster.

The main concept of LIT is a structured process for assessing congruence between clusters and morphospecies. For this purpose, LIT makes suggestions which specimens should be studied for cluster validation, and how conflict between molecular and morphological evidence should be resolved. The LIT rules are based on an empirically study that evaluated several potential predictors of conflict between molecular clusters and morphospecies: (1) cluster stability across distance thresholds, (2) maximum intracluster genetic distance, (3) number of haplotypes, (4) number of specimens in cluster, and (5) measures of geographic representation. The analysis revealed that incongruence between barcode clusters and morphology was best predicted by cluster stability and maximum pairwise distance. Clusters that were stable across thresholds turned out to have a low risk of being incongruent and are according to the LIT rules sampled minimally, by examining a pair of specimens representing the most divergent haplotypes. In contrast, clusters whose boundaries shifted across thresholds required closer scrutiny. The most effective strategy for detecting incongruence was examining representatives of three haplotypes, the most abundant haplotype and the ones yielding the maximum pairwise distances within a cluster. This approach detected most cases of conflict, with remaining cases found by applying the same rules also to stable clusters with an intracluster variability of >1.5%. As a default, IntegraTax uses 1-3% as an instability zone and 1.5% as the variability threshold in classifying clusters as either stable or instable. These settings can be changed and specimen selection and classifications get dynamically updated.

LIT treats DNA-based clusters as a hierarchy of nested species hypotheses and uses morphology to test which hypothesis is in agreement with observed phenotypic variation. Three outcomes are possible when the congruence between cluster and morphology is assessed. First, congruence, where morphology is in agreement with the specimens in the focal cluster. Second, incongruence due to lumping, where morphology suggests two or more species within the focal cluster, corresponding to groupings at lower levels of the hierarchy. Third, incongruence with splitting, where morphology suggests a single species, but one that is identical to specimens from the next higher, more inclusive cluster in the hierarchy. This indicates that the genetic structure within the cluster is intra-rather than interspecific. Because clusters are hierarchical, testing all clusters for agreement with morphology would be impractical. Instead, analyses begin at a clustering threshold expected to show high overall agreement with morphology. If the threshold is set too low, many clusters will disagree with morphology and need to be fused, increasing the workload because more clusters must be examined. If the threshold is set too high, many clusters will need to be subdivided, increasing the workload because more cross-cluster comparisons are required. However, the outcome will in principle be the same regardless of the initial threshold chosen. This is because morphology will ultimately settle on the same clusters to be congruent. Any observed differences when LIT is applied at different thresholds could only arise from alternative haplotype-testing paths taken to reach that level. Note that LIT only considers sequences and morphology to be genuinely incongruent, when none of the molecular hypotheses align with morphology. This is when additional data are required to resolve the species limits.

#### Implementation

The taxonomy module operates on an output file with an *.itv extension produced by the clustering module. It is opened in “*IntegraTaxViz*.*html*,” a browser-based visualization environment written in JavaScript using the D3 library. The taxonomic work is carried out in this interactive environment by interpreting the hierarchical barcode cluster diagram and associated cluster statistics generated during clustering. The taxonomy module does not generate new clusters or infer species boundaries but operates exclusively on existing clustering results. For ease of use, the name and location of the *.itv file is displayed once *IntegraTaxViz*.*html* is opened after clustering. *IntegraTaxViz* also provides an export panel that allows users to export SVG, FASTA, Excel LIT only, and SPART files directly from the interface.

*IntegraTaxViz* enables interactive exploration of the hierarchical barcode clustering, allowing users to navigate cluster structure and inspect genetic distances at different levels of the hierarchy. Context menus provide annotation options. Some are shared between specimen and internal nodes. They include (1) entering or editing species names, (2) attaching notes, and (3) the option to lock decisions. Additional specimen-level annotation options include marking specimens as examined or damaged and recording sex or the availability of a slide-mount. In addition, the barcode of a specimen can be copied for a BLAST online. Node-level annotation options include collapsing and expanding of subtrees to reduce the size of the visible cluster fusion diagram. All these capabilities are designed to support integrative taxonomy projects at scale, where manual tracking of decisions across large numbers of specimens would otherwise become impractical. FASTA export is also available both at node- and project-wide level. The sequence headers then include the annotations and species names. Node-level FASTA exports also allow for curated subsets of sequences to be forwarded directly to external tools such as BLAST (Altschul et al., 1990), BOLD (Ratnasingham et al., 2024), BOLD View (https://bold-view-bf2dfe9b0db3.herokuapp.com/), or GBIF sequence identification (https://www.gbif.org/tools/sequence-id).

Within the interactive *IntegraTaxViz* environment, the taxonomy module provides operational support for evaluating molecular clusters and selecting specimens for examination following the LIT framework described above. Clusters are visually flagged as “PI” (red branches, Figure 2) or “non-PI” (blue branches), where PI denotes unstable, potentially incongruent clusters sensu Hartop et al. (2022), based on an instability zone that is defined by upper and lower instability bound set by the user via a menu. The same menu also allows for setting a maximum intracluster p-distance that triggers more extensive intra-cluster sampling. These settings implement the empirically derived LIT predictors into the software while leaving all taxonomic decisions to the user. In addition, a summary function allows users to track project progress by counting specimens with assigned species names and to calculate clustering-based metrics such as match ratio and specimen congruence across all possible thresholds, which can be used to guide threshold choice in datasets with valid species names. Should users desire to not use recommendations by IntegraTax, they can import species delimitations obtained by other software in SPART format, assess areas of dendrograms that show variations across partitioning schemes and use the annotation features of IntegraTax to proceed with taxonomic work.

**Figure 2:**
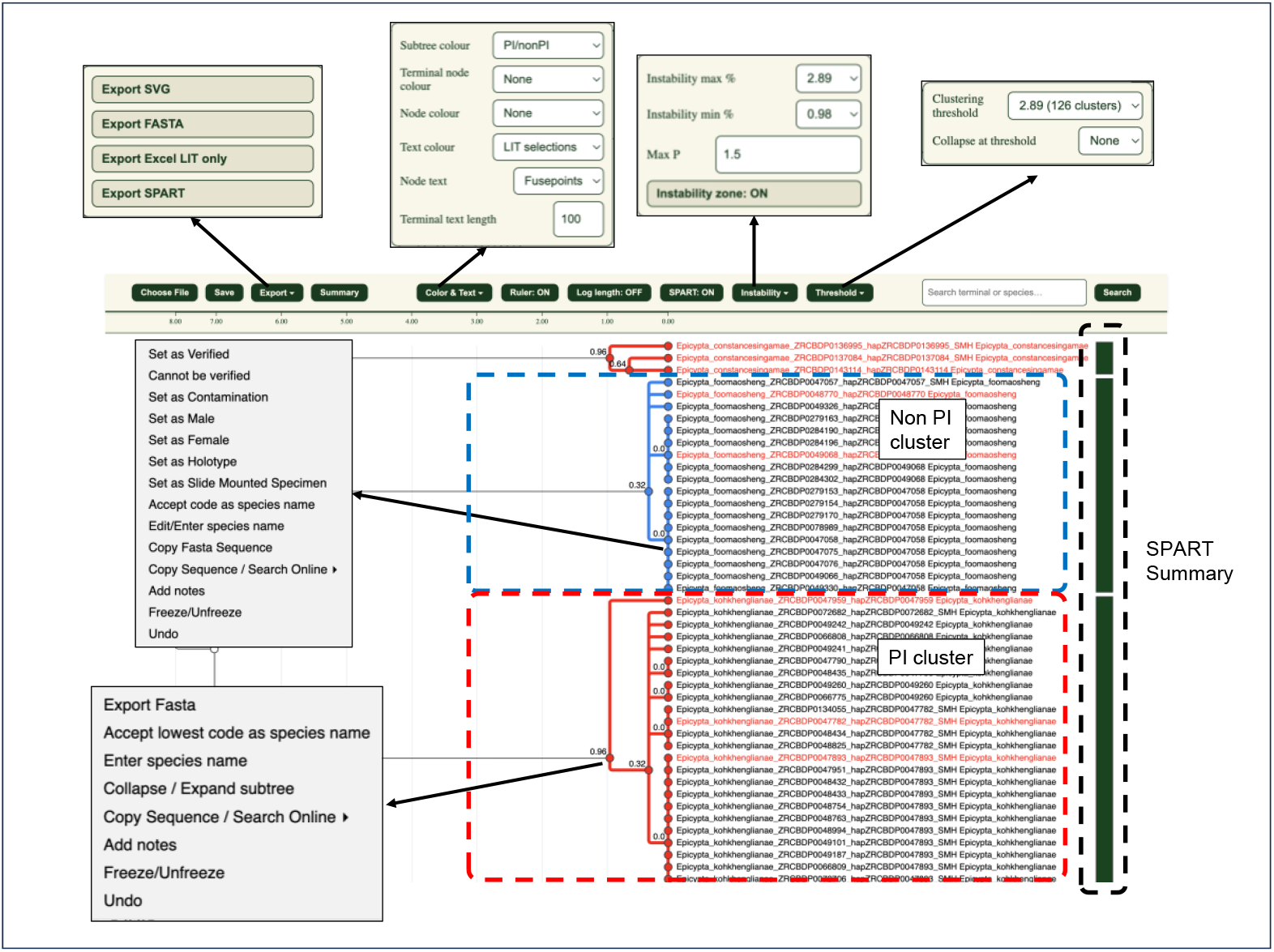
A screenshot of *IntegraTaxViz*.*html* with dialogs which allow you to visualize and annotate an integrative taxonomy project. Red and blue clusters highlight the categorizations into PI and non-PI clusters. The specimens in red are the recommendations made by IntegraTax for specimen selection to obtain a secondary data source. The green bar on the right is the summary of the species delimitation schemes in the SPART output. Tool tips on this bar will inform the number of partitioning schemes, while split bars will appear if partitioning schemes are in disagreement.

### Empirical test of IntegraTax using Singapore Mycetophilidae

We evaluated IntegraTax using Dataset 1 of Singapore Mycetophilidae (Meier et al., 2025b), comprising 1,454 barcoded specimens, to assess the robustness of the LIT workflow to parameter choice and the concordance between barcode clusters and species assignments.

#### Robustness of LIT workflow

We assessed how specimen recommendations produced by IntegraTax vary with changes in the instability range and maximum pairwise distance (maxP) parameters. When maxP was fixed at 1.5%, increasing the instability range from 1-2% to 1-5% resulted in only a minor reduction in the number of specimens recommended for examination (from 213 to 209 specimens, corresponding to 14.65% to 14.37% of the dataset; Table 1). Similarly, when the instability range was fixed at 1-3%, increasing maxP from 1% to 3% reduced the number of recommended specimens from 222 to 202 (15.27% to 13.89% of specimens).

**Table 1.** Number of specimens recommended by IntegraTax as instability range and maxP parameters are varied.

| Scenario | Parameter varied | Range/Value tested | # Specimens recommended | % of specimens |
| --- | --- | --- | --- | --- |
| Fixed maxP = 1.5% | Instability range | 1-2% | 213 | 14.65% |
|  |  | 1-3% | 212 | 14.58% |
|  |  | 1-4% | 211 | 14.51% |
|  |  | 1-5% | 209 | 14.37% |
| Fixed instability = 1–3% | maxP | 1% | 222 | 15.27% |
|  |  | 1.5% | 212 | 14.58% |
|  |  | 2% | 205 | 14.10% |
|  |  | 2.5% | 205 | 14.10% |
|  |  | 3% | 202 | 13.89% |

Across all tested parameter combinations, IntegraTax consistently reduced the taxonomic workload to the examination of approximately 15% of specimens, with changes of <1.4%, indicating robustness to parameter variation. In cases where changes in clustering thresholds grouped specimens differently, the exact set of specimens selected for examination vary leading to the variations in number of specimens recommended in Table 1; however, these differences yielded the same final taxonomic conclusion, as specimens within nested or alternative clustering schemes belonged to the same species (see Figure 3).

**Figure 3:**
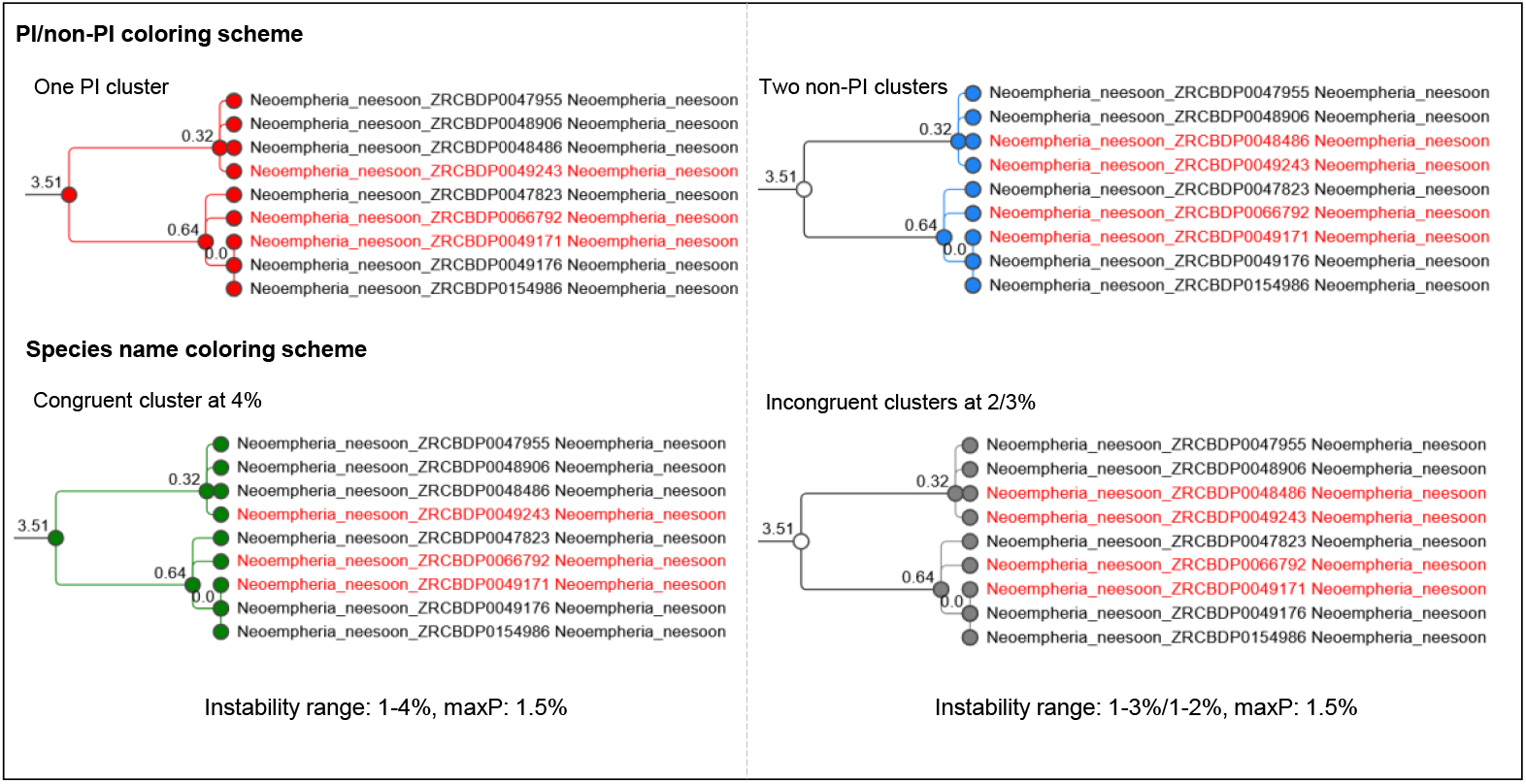
Visualization of specimen grouping in IntegraTax. Specimens belonging to a single species are grouped differently under alternative clustering/instability parameter settings. PI and non-PI clusters are shown in red and blue in top panel, respectively, and species-based coloring is shown below (green = congruent, red = incongruent). Changing the instability range (1-4% versus 1-3%/1-2% at maxP = 1.5%) alters PI/non-PI assignment and the number of specimens recommended for examination (three versus four).

#### Threshold choice and congruence with morphology

Across distance thresholds from approximately 2% to 5%, the match ratio increased from 0.873 to 0.974, while specimen congruence increased from 0.844 to 0.922 (Table 2). The number of congruent clusters and specimens in congruent clusters increased at higher thresholds and this dataset reveals deeper divergences in COI within species.

**Table 2:** Summary statistics for Singapore Mycetophilidae dataset 1.

| Summary statistics | 2% (1.93%) | 3% (2.89%) | 4% (3.88%) | 5% (4.85%) |
| --- | --- | --- | --- | --- |
| Match ratio | 0.873 | 0.913 | 0.95 | 0.974 |
| Clusters with species names | 128 | 124 | 121 | 115 |
| Congruent clusters | 107 | 110 | 113 | 113 |
| Specimen congruence | 0.844 | 0.851 | 0.904 | 0.922 |
| Specimens in congruent clusters | 1227 | 1237 | 1314 | 1340 |
| Specimens with species names | 1454 | 1454 | 1454 | 1454 |

## Discussion

The detailed taxon-specific knowledge of experts is the most valuable resource in taxonomy. Taxonomists deserve tools that enhance the efficiency and accessibility of taxonomic work, without being forced to learn unnecessarily complex pipelines and/or multiple software packages to carry out their work. Sequence clustering methods should clearly visualize the underlying genetic data through transparent and reproducible procedures, and taxonomic decisions should be easy to record, inspect, and revisit within a coherent working environment. At the same time, results should be exportable and importable in formats suitable for publication and for further exploration using complementary tools. While existing software solutions provide individual components of such a pipeline (see tools in iTaxoTools (Vences et al., 2021)), these elements are rarely integrated in a way that supports the full process of taxonomic reasoning, from data exploration to documented decisions. IntegraTax was developed to bring these processes together by linking deterministic clustering, interactive annotation, and flexible export, thereby supporting integrative taxonomy as a continuous and well-documented practice.

An integrated software package is needed because the widespread adoption of integrative taxonomy depends on the accessibility and autonomy of the supporting infrastructure. Recent advances in sequencing technologies allow DNA barcoding to be carried out efficiently beyond a small number of centralized facilities (Jabot et al., 2025, Sánchez-Vendizú et al., 2025, Zomoroka and Zinenko, 2025). Scientists in species-rich regions can now participate more easily in biodiversity research, but this shift toward decentralized workflows also places new requirements on analytical tools. They must facilitate the inspection of data, documentation of decisions, and reproduce results independently of proprietary platforms or opaque processing steps. This need is heightened by the fact that centralized barcoding infrastructures have often relied on processing steps and identifiers that are difficult to reproduce or scrutinize independently (Meier et al., 2022). As biodiversity is distributed unevenly across the globe, the ability to study it must be similarly distributed. IntegraTax tries to overcome these limitations by being an open-access, reusable environment that provides analytical autonomy, allowing for reproducible taxonomic work across geographically distributed research groups. As the *itv file is storing the annotation information, and only a browser is needed to view it, it facilitates collaborative taxonomic work.

IntegraTax operates firmly within the conceptual framework of integrative taxonomy in which sequence data are used to generate provisional species hypotheses rather than being used to delimit species directly. For most multicellular taxa, morphology, geographic information, and ecological evidence are available, making exclusive reliance on a single character system undesirable and unnecessary. This is why we would argue that sequence data should mostly be used for structuring large specimen sets into putative species that are then evaluated using other evidence. IntegraTax uses single-linkage clustering, but the taxonomy module is compatible with the outputs of species delimitation algorithms such as ASAP, ABGD, and phylogeny-based approaches. Note, however, that IntegraTax was designed primarily for protein-encoding, largely intron-free markers such as animal *COI* barcodes, which are widely used and have well-understood properties. The framework can be used for other sequence data, including nuclear markers and multi-locus datasets. However, extending its use beyond *COI* requires careful consideration of marker-specific properties, including differences in evolutionary rates, sequence length variation, and patterns of divergence across loci or genomic regions.

Species delimitation is not an endpoint in taxonomy, but only a step toward species identification, diagnosis, and formal description. This distinction is important because, despite the widespread use of molecular data for species delimitation, molecular diagnoses remain rare in species descriptions, particularly for insects, which comprise most of the undescribed animal diversity. Although IntegraTax is primarily a tool for delimiting species hypotheses, it also facilitates species identification when reliably identified reference sequences are available from external databases. In such cases, clustering can directly assign a species name to a specimen cluster. IntegraTax can also contribute directly to species descriptions because molecular diagnoses can be computed based on exporting FASTA datasets as long as species names are in the FASTA headers. This input is then suitable for UITOTO (Torres et al., 2025) which is designed to generate contrastive molecular diagnoses. IntegraTax can, however, not help with resolving taxonomic names of unclear or unstable meaning.

Taken together, IntegraTax is an extensible framework rather than a closed solution. While the current implementation focuses on clustering and hypothesis tracking based on sequence data, the same structure can accommodate additional information such as images, geographic data, ecological traits, or other specimen-level metadata. As IntegraTax is implemented as a browser-based environment, it can also be linked to external web applications, allowing functionality to expand without constraining workflows to a fixed software ecosystem. We would argue that this flexibility is essential for the future of integrative taxonomy, where the types of evidence used to evaluate species boundaries will vary across taxa and study systems. By emphasizing transparent workflows, explicit hypothesis testing, and interoperability with complementary tools, IntegraTax will thus hopefully provide a solid foundation for small- and large-scale integrative taxonomy projects.

## Acknowledgements

We are grateful to Vladimir Blagoderov, Pierfilippo Cerretti, Olavi Kurina, Jostein Kjærandsen, Aleida Ascenzi, Ronniel Pedales, Bernardo Santos, Dalton de Souza Amorim, Ivan Neo and Daniel Gustafsson for testing IntegraTax, identifying the bugs, supporting and encouraging its development.

## References

Ahrens, D., Fujisawa, T., Krammer, H. J., Eberle, J., Fabrizi, S. & Vogler, A. P. 2016. Rarity and incomplete sampling in DNA-based species delimitation. Systematic Biology, 65, 478–494.

Altschul, S. F., Gish, W., Miller, W., Myers, E. W. & Lipman, D. J. 1990. Basic local alignment search tool. J Mol Biol, 215, 403–10.

Amorim, D. D. S., Oliveira, S. S., Balbi, M. I. P. A., Ang, Y., Torres, A., Yeo, D., Srivathsan, A. & Meier, R. 2025. An integrative taxonomic treatment of the Mycetophilidae (Diptera: Bibionomorpha) from Singapore reveals 115 new species on 730km^2^. bioRxiv.

Bánki, O., Roskov, Y., Döring, M., Ower, G., Hernández Robles, D.R., Plata Corredor, C.A., Stjernegaard Jeppesen, T., Örn, A., Pape, T., Hobern, D., Garnett, S., Little, H., Dewalt, R. E., Miller, J., Orrell, T., Aalbu, R., Abbott, J., Abreu, C. & P.A., A. 2025. Catalogue of Life (2025-11-16 XR). Catalogue of Life Foundation, Amsterdam, Netherlands.

Blaxter, M., Mann, J., Chapman, T., Thomas, F., Whitton, C., Floyd, R. & Abebe, E. 2005. Defining operational taxonomic units using DNA barcode data. Philosophical Transactions of the Royal Society B-Biological Sciences, 360, 1935–1943.

Caruso, V., Hartop, E., Chimeno, C., Noori, S., Srivathsan, A., Haas, M., Lee, L., Meier, R. & Whitmore, D. 2024. An integrative framework for dark taxa biodiversity assessment at scale: A case study using (Diptera, Phoridae). Insect Conservation and Diversity, 17, 968–987.

Collins, R. A., Boykin, L. M., Cruickshank, R. H. & Armstrong, K. F. 2012. Barcoding’s next top model: an evaluation of nucleotide substitution models for specimen identification. Methods in Ecology and Evolution, 3, 457–465.

Collins, R. A. & Cruickshank, R. H. 2013. The seven deadly sins of DNA barcoding. Molecular Ecology Resources, 13, 969–975.

Daily, J. 2016. Parasail: SIMD C library for global, semi-global, and local pairwise sequence alignments. Bmc Bioinformatics, 16.

Dayrat, B. 2005. Towards integrative taxonomy. Biological Journal of the Linnean Society, 85, 407–415.

Fujisawa, T. & Barraclough, T. G. 2013. Delimiting species using single-locus data and the Generalized Mixed Yule Coalescent approach: A revised method and evaluation on simulated data sets. Systematic Biology, 62, 707–724.

GBIF. 2025. About species counts in GBIF [Online]. Available: https://www.gbif.org/about-species-counts [Accessed 15.12.2025 2025].

Gustafsson, D. R., Lee, L., Grossi, A. A., Zou, F., Tan, D. J. X., Hwang, W. S. & Meier, R. 2025. From host to host, and continent to continent: two phoresy-enabled Guimaraesiella hitchhiker louse species revealed by integrative taxonomy (Phthiraptera: Ischnocera). . Medical and Veterinary Entomology, In Press.

Hartop, E., Srivathsan, A., Ronquist, F. & Meier, R. 2022. Towards large-scale integrative taxonomy (LIT): Resolving the data conundrum for dark taxa. Systematic Biology, 71, 1404–1422.

Hawlitschek, O., Nagy, Z. T., Berger, J. & Glaw, F. 2013. Reliable DNA barcoding performance proved for species and island populations of Comoran squamate reptiles. Plos One, 8.

HÉBert, C. & Favret, C. 2025. Large-scale integrative taxonomy of the smallest insects reveals astonishing temperate diversity (Hymenoptera: Chalcidoidea: Mymaridae). Molecular Ecology.

Hebert, P. D. N., Braukmann, T. W. A., Prosser, S. W. J., Ratnasingham, S., Dewaard, J. R., Ivanova, N. V., Janzen, D. H., Hallwachs, W., Naik, S., Sones, J. E. & Zakharov, E. V. 2018. A Sequel to Sanger: amplicon sequencing that scales. Bmc Genomics, 19.

Jabot, F., Auger, G., Bonnal, P., Pizaine, M., Roncoroni, M., Revaillot, S. & Pottier, J. 2025. Use of massive DNA barcoding to monitor biodiversity: A test on forest soil macrofauna. Forest Ecology and Management, 595.

Lee, L., OboŇA, J., Lee, Y. X., Lee, K. T., Puniamoorthy, J., Tan, L. Y. K., Choo, R., Tan, D. J. X., Kwak, M., Ang, Y. & Meier, R. 2025. Avian afterlives: Integrative taxonomy of hippoboscid flies (Diptera) from citizen-reported bird carcasses reveals a new species and host–parasite diversity in Singapore. Journal of Zoological Systematics and Evolutionary Research, 2025.

Meier, R., Blaimer, B. B., Buenaventura, E., Hartop, E., Rintelen, T., Srivathsan, A. & Yeo, D. 2022. A re-analysis of the data in Sharkey et al.’s (2021) minimalist revision reveals that BINs do not deserve names, but BOLD Systems needs a stronger commitment to open science. Cladistics, 38, 264–275.

Meier, R., Lawniczak, M. K. N. & Srivathsan, A. 2025a. Illuminating entomological dark matter with DNA barcodes in an era of insect decline, deep learning, and genomics. Annual Review of Entomology, 70, 185–204.

Meier, R., Shiyang, K., Vaidya, G. & Ng, P. K. L. 2006. DNA barcoding and taxonomy in diptera: A tale of high intraspecific variability and low identification success. Systematic Biology, 55, 715–728.

Meier, R., Srivathsan, A., Oliveira, S. S., Balbi, M. I. P. A., Ang, Y., Yeo, D. R., Kjaerandsen, J. & Amorim, D. D. 2025b. “Dark taxonomy”: A new protocol for overcoming the taxonomic impediments for dark taxa and broadening the taxon base for biodiversity assessment. Cladistics, 41, 223–238.

Meier, R., Wong, W. H., Srivathsan, A. & Foo, M. S. 2016. $1 DNA barcodes for reconstructing complex phenomes and finding rare species in specimen-rich samples. Cladistics, 32, 100–110.

Miralles, A., Ducasse, J., Brouillet, S., Flouri, T., Fujisawa, T., Kapli, P., Knowles, L. L., Kumari, S., Stamatakis, A., Sukumaran, J., Lutteropp, S., Vences, M. & Puillandre, N. 2022. SPART: A versatile and standardized data exchange format for species partition information. Molecular Ecology Resources, 22, 430–438.

Mora, C., Tittensor, D. P., Adl, S., Simpson, A. G. B. & Worm, B. 2011. How many species are there on earth and in the ocean? Plos Biology, 9, e1001127.

Padial, J. M., Miralles, A., DE LA Riva, I. & Vences, M. 2010. The integrative future of taxonomy. Frontiers in Zoology, 7.

Page, R. 2025. Tracking changes in DNA barcode BINs [Online]. Available: https://iphylo.blogspot.com/2025/05/tracking-changes-in-dna-barcode-bins.html [Accessed 15.12.2025 2025].

Pentinsaari, M., Salmela, H., Mutanen, M. & Roslin, T. 2016. Molecular evolution of a widely-adopted taxonomic marker (COI) across the animal tree of life. Scientific Reports, 6.

Pons, J., Barraclough, T. G., Gomez-Zurita, J., Cardoso, A., Duran, D. P., Hazell, S., Kamoun, S., Sumlin, W. D. & Vogler, A. P. 2006. Sequence-based species delimitation for the DNA taxonomy of undescribed insects. Systematic Biology, 55, 595–609.

Ramirez, J. L., Valdivia, P., Rosas-Puchuri, U. & Valdivia, N. L. 2023. SPdel: A pipeline to compare and visualize species delimitation methods for single-locus datasets. Molecular Ecology Resources, 23, 1959–1965.

Ratnasingham, S., Wei, C., Chan, D., Agda, J., Ballesteros-Mejia, L., Ait Boutou, H., El Bastami, Z. M., Ma, E., Manjunath, R., Rea, D., Ho, C., Telfer, A., Mckeowan, J., Rahulan, M., Steinke, C., Dorsheimer, J.M. M. & Hebert, P. D. N. 2024. BOLD v4: A centralized bioinformatics platform for DNA-based biodiversity data. In: Salle, R. D. (ed.) DNA Barcoding: Methods and Protocols. New York: Springer US.

Riccardi, P. R. & Hartop, E. 2024. Large-scale integrative taxonomy of Swedish grass flies (Diptera, Chloropidae) reveals hitherto unknown complexity of a dark taxon. Zoologica Scripta, 53, 614–631.

SÁNchez-VendizÚ, P., Erkenswick, G., Reyes, J., Clinton, S. L., Espejo, T. S., CÁCeres, G., Libke, Z., Arana, A., Mendoza-Silva, J., Tirapelle, C., Williams, S., Swamy, V., Martínez-Altamirano, J., Esteves, J., Barnuevo-Bullón, J. P., Hernández-Mejía, J., Caffo, X., Malpica, A. M., Salazar-AragóN, R., GutiÉRrez-JimÉNez, L., Stabile, J., Cuzmar, N., Paine, T. D., Peralta-Aguilar, P., Inga-Díaz, G., Lescano, J., Viñas-Martínez, A., Mcelroy, M. E., Coayla, D., Linares, R. L. M., Pilfold, N. W., Sacco, A. J., Arakaki, M., Mena, J. L., Tobler, M. W., Salinas, L., Arana, C., Pacheco, V., Prost, S. & Watsa, M. 2025. Decoding the Peruvian Amazon with DNA barcoding of vertebrate and plant taxa. Scientific Data, 12.

Santos, B. F., Srivathsan, A., Neves, K. & Meier, R. 2025. Weak and inverse latitudinal diversity gradients in the 10 most abundant and diverse flying insect clades. bioRxiv.

Slater-Baker, M. R., Fagan-Jeffries, E. P., Oestmann, K. J., Portmann, O. G., Bament, T. M., Howe, A. G., Guzik, M. T., Bradford, T. M., Mcclelland, A. R., Woodward, A., Clarke, S., Ducker, N. & FernÁNdez-Triana, J. 2025. DNA barcoding, integrative taxonomy, citizen science, and Bush Blitz surveys combine to reveal 34 new species of Apanteles (Hymenoptera, Braconidae, Microgastrinae) in Australia. Zookeys, 1–128.

Sosic, M. & Sikic, M. 2017. Edlib: a C/C plus plus library for fast, exact sequence alignment using edit distance. Bioinformatics, 33, 1394–1395.

Srivathsan, A., Ang, Y., Heraty, J. M., Hwang, W. S., Jusoh, W. F. A., Kutty, S. N., Puniamoorthy, J., Yeo, D., Roslin, T. & Meier, R. 2023. Convergence of dominance and neglect in flying insect diversity. Nat Ecol Evol, 7, 1012–1021.

Srivathsan, A., Hartop, E., Puniamoorthy, J., Lee, W. T., Kutty, S. N., Kurina, O. & Meier, R. 2019. Rapid, large-scale species discovery in hyperdiverse taxa using 1D MinION sequencing. Bmc Biology, 17.

Srivathsan, A., Lee, L., Katoh, K., Hartop, E., Kutty, S. N., Wong, J., Yeo, D. & Meier, R. 2021. ONTbarcoder and MinION barcodes aid biodiversity discovery and identification by everyone, for everyone. BMC Biol, 19, 217.

Srivathsan, A. & Meier, R. 2012. On the inappropriate use of Kimura-2-parameter (K2P) divergences in the DNA-barcoding literature. Cladistics, 28, 190–194.

Takezaki, N. 1998. Tie trees generated by distance methods of phylogenetic reconstruction. Molecular Biology and Evolution, 15, 727–737.

Torres, A., Lee, L., Srivathsan, A. & Meier, R. 2025. UITOTO: a software for generating molecular diagnoses for species descriptions. bioRxiv.

Vences, M., Miralles, A., Brouillet, S., Ducasse, J., Fedosov, A., Kharchev, V., Kostadinov, I., Kumari, S., Patmanidis, S., Scherz, M. D., Puillandre, N. & Renner, S. S. 2021. iTaxoTools 0.1: Kickstarting a specimen-based software toolkit for taxonomists. Megataxa, 6.

Vogel, J., Forshage, M., Bartsch, S. B., Ankermann, A., Mayer, C., Von Falkenhausen, P., Rduch, V., Mueller, B., Braun, C., Krammer, H. J. & Peters, R. S. 2024. Integrative characterisation of the Northwestern European species of Dalman, 1823 (Hymenoptera, Cynipoidea, Figitidae) with the description of three new species. Journal of Hymenoptera Research, 97, 621–698.

Wang, W. Y., Srivathsan, A., Foo, M., Yamane, S. K. & Meier, R. 2018. Sorting specimen-rich invertebrate samples with cost-effective NGS barcodes: Validating a reverse workflow for specimen processing. Molecular Ecology Resources, 18, 490–501.

Will, K. W. & Rubinoff, D. 2004. Myth of the molecule: DNA barcodes for species cannot replace morphology for identification and classification. Cladistics, 20, 47–55.

Yeo, D., Srivathsan, A. & Meier, R. 2020. Longer is not always better: Optimizing barcode length for large-scale species discovery and identification. Syst Biol, 69, 999–1015.

Zhang, J. J., Kapli, P., Pavlidis, P. & Stamatakis, A. 2013. A general species delimitation method with applications to phylogenetic placements. Bioinformatics, 29, 2869–2876.

Zomoroka, A. & Zinenko, O. 2025. Molecular cross-validation of intraspecific structure of Rutpela maculata (Insecta: Coleoptera: Cerambycidae). Journal of Vasyl Stefanyk Precarpathian National University, 12, 46–59.

